# Tissue-Specific Ablation of Liver Fatty Acid-Binding Protein Induces a Metabolically Healthy Obese Phenotype in Female Mice

**DOI:** 10.1101/2025.01.02.631082

**Authors:** Hiba Radhwan Tawfeeq, Atreju I Lackey, Yinxiu Zhou, Anastasia Diolointzi, Sophia Zacharisen, Yin Hei Lau, Loredana Quadro, Judith Storch

## Abstract

**Background/Objectives:** Obesity is associated with numerous metabolic complications including insulin resistance, dyslipidemia, and a reduced capacity for physical activity. Whole-body ablation of liver fatty acid-binding protein (LFABP) in mice was shown to alleviate several of these metabolic complications; high fat (HF) fed LFABP knockout (LFABP^-/-^) mice developed higher fat mass than their wild-type (WT) counterparts but displayed a metabolically healthy obese (MHO) phenotype with normoglycemia, normoinsulinemia, and reduced hepatic steatosis compared with WT. LFABP is expressed in both liver and intestine, thus in the present study, LFABP conditional knockout (cKO) mice were generated to determine the contributions of LFABP specifically within the liver or the intestine to the whole body phenotype of the global knockout.

**Methods:** Female liver-specific LFABP knockout (LFABP^liv-/-^), intestine-specific LFABP knockout (LFABP^int-/-^), and floxed LFABP (LFABP^fl/fl^) control mice were fed a 45% Kcal fat semipurified HF diet for 12 weeks.

**Results:** While not as dramatic as found for whole-body LFABP^-/-^ mice, both LFABP^liv-/-^ and LFABP^int-/-^ mice had significantly higher body weights and fat mass compared with LFABP^fl/fl^ control mice. As with the global LFABP nulls, both LFABP^liv-/-^ and LFABP^int-/-^ mice remained normoglycemic and normoinsulinemic. Despite their greater fat mass, the LFABP^liv-/-^ mice did not develop hepatic steatosis. Additionally, LFABP^liv-/-^ and LFABP^int-/-^ mice had higher endurance exercise capacity when compared with LFABP^fl/fl^ control mice.

**Conclusions:** The results suggest, therefore, that either liver-specific or intestine-specific ablation of LFABP in female mice is sufficient to induce, at least in part, the MHO phenotype observed following whole-body ablation of LFABP.

## 1. Introduction

Obesity, a disease state that is often accompanied by an array of metabolic comorbidities, is one of the leading causes of preventable death globally [1]. With increased fat storage being integral to the development of obese phenotypes, dietary lipid quantity and quality are thought to play important roles in the etiology of obesity and its related health effects [2–4]. As such, understanding how metabolically active tissues, such as the small intestine (SI) and liver, handle exposure to endogenous and exogenous lipid is important in determining or updating guidance associated with reducing the medical, quality of life, and economic burdens of obesity.

The absorptive enterocytes of the SI are primarily responsible for the uptake and subsequent processing and delivery of the products of dietary lipid digestion, primarily fatty acids (FAs) and monoacylglycerols (MGs), to the periphery [5], while the liver plays a major role in importing, synthesizing, storing, and exporting lipids [6,7]. Liver fatty acid-binding protein (LFABP or FABP1) is an intracellular protein that is abundantly expressed in both SI epithelial cells, and in liver hepatocytes and hepatic stellate cells [8–12]. Previous *in vitro* studies have demonstrated that LFABP binds FA at high affinity, with K_d_ values in the nanomolar range [13,14]. Additionally, LFABP also binds other types of lipids, including but not limited to MGs, prostaglandins, lysophospholipids, endocannabinoids, and cholesterol [15–20].

We found that male whole-body LFABP^-/-^ mice had increased body weight gain and fat mass (FM) accumulation in response to chronic high fat (HF) feeding [21]. This obese phenotype may be caused, in part, by the higher food intake and the slower intestinal transit time of LFABP^-/-^ relative to WT mice [21–23]. Although these mice become obese they appear to be relatively healthy, remaining normoglycemic and normoinsulinemic, having reduced hepatic steatosis, and having intestinal triglyceride (TG) secretion rates similar to those of lean mice [21,24–26]. Further, despite their obesity, the LFABP^-/-^ mice are more active [21] and have greater exercise endurance than WT mice [27]. Fecal short chain FA (SCFA) levels were also found to be higher in LFABP^-/-^ mice when compared with WT mice, which may partly explain some of the beneficial metabolic changes that are observed in LFABP^-/-^ mice [22].

It has been recognized that a subset of the obese population is nevertheless healthy, not displaying various comorbidities that are commonplace amongst obese people; this phenomenon has become known as the “metabolically healthy but obese” (MHO) state [28–31]. The LFABP^-/-^ mice, thus, appear to be a model of MHO. To understand the underlying causes of the LFABP^-/-^ phenotype, it is critical to know whether it is dependent on the ablation of LFABP in the liver, the intestine, or both the liver and the intestine.

Biological sex appears to influence LFABP expression, with female rats having increased liver LFABP expression when compared with male rats [32,33]. Interestingly, estradiol treatment of castrated male rats resulted in hepatic LFABP levels similar to those of intact female rats, while testosterone treatment of ovariectomized female rats resulted in hepatic LFABP levels similar to those of intact males, showing that sex steroids play a role in the regulation of LFABP expression [34]. We have recently found that, similar to males, female LFABP^-/-^ mice fed a HF diet (HFD) also gain significantly more weight and FM, when compared with their WT counterparts [35].

To determine the role of LFABP specifically within the liver or the intestine, in the present studies we report the generation of 2 lines of conditional knockout (cKO) mice in which the LFABP gene is ablated solely in liver or solely in intestine. Data from female mice demonstrate the overlapping contributions of liver-LFABP and intestinal-LFABP to the MHO phenotype observed in the whole-body LFABP null mice.

## 2. Materials and Methods

### Generation of LFABP Floxed Mice

LFABP floxed mice (LFABP^fl/fl^) were generated at the Rutgers Genome Editing Core Facility using clustered regulatory interspaced short palindromic repeats (CRISPR)/CRISPR-associated Cas (Cas9) protein technology to introduce 2 loxP sites flanking exons 2 and 3 of the gene encoding LFABP (More details can be found in the supplemental material).

### Generation of conditional LFABP Null Mice

The LFABP^fl/fl^ mice were bred with mice that were either homozygous for Cre recombinase driven by the albumin promoter (A-cre), or heterozygous for Cre recombinase driven by the viliin promoter (V-cre; The Jackson Laboratory), to generate double-mutants (LFABP^fl/+,Acre/+^; LFABP^fl/+,Vcre/+^ respectively). These mice were then backcrossed with the control WT (LFABP^fl/fl^) to generate either liver-specific LFABP KO (LFABP^liv-/-^) or intestine-specific LFABP KO (LFABP^int-/-^) and littermate LFABP^fl/fl^ mice. Mice were maintained on a 12-hour light/dark cycle, and a controlled temperature. They were allowed *ad libitum* access to standard rodent chow (Purina Laboratory Rodent Diet 5015) until the start of the HF feeding period at 2 months of age.

### DNA Extraction for Genotyping

DNA extraction was performed as described previously [36]. For the genotyping of the LFABP^fl/fl^ mice, 4 primers were developed and used to assess the upstream and downstream loxP sites in 2 separate PCR reactions. The primer sequences for the LFABP^fl/fl^ protocols were as follows:

1- Primers used for the upstream loxP:

FABP1A: 5’-AGACAAGTCAAAGATCATGAATGTGAG-3’

FABP1B: 5’-TGGCTCTTAGAGTGGGAACACTTC-3’

2- Primers used for the downstream loxP: FABP1C: 5’-CGGAGTTGATAGATATCAGATC-3’

FABP1D: 5’-GAAACAGGGCAAGGCCAGCTATG-3’

After the reactions, PCR products for the upstream loxP reaction were digested with PsiI, while the PCR products for the downstream loxP site were digested with EcoRI. Then, electrophoresis was performed on a 2% agarose gel. WT mice that did not have the inserted loxP sites only had 1 band for both the upstream (320 BP) and downstream (506 BP) reactions, while LFABP^fl/fl^ mice had 2 smaller bands for both the upstream (231 BP and 119 BP) and downstream (325 BP and 215 BP) reactions.

The genotyping protocol for the Acre mice used 3 primers for 1 PCR reaction. One primer, Acre common, was shared with both WT and Mutant primers. The Acre WT primer was used to detect a 351 BP band in WT mice, while the Acre mutant primer was used to detect a 390 BP band in Acre mice. The Acre mice could be hemizygotes or homozygotes. The primer sequences used for the Acre genotyping protocol were as follows:

Acre reaction (WT and mutant bands):

Acre Common: 5’-TTG GCC CCT TAC CAT AAC TG-3’

Acre WT: 5’-TGC AAA CAT CAC ATG CAC AC-3’

Acre Mutant: 5’-GAA GCA GAA GCT TAG GAA GAT GG-3’

The genotyping protocol for the Vcre mice used 3 primers for 2 separate PCR reactions. One primer, Vcre common, was used for both reactions. The Vcre WT primer was used to detect a 186 BP band in WT mice, while the Vcre mutant primer was used to detect a 150 BP band in Vcre/+ mice. Since the Vcre/+ mice must be maintained as hemizygotes, bands for both the Vcre WT reaction and the Vcre mutant reaction would be present [37]. The primer sequences for the Vcre genotyping protocols were as follows:

Vcre WT reaction

Vcre Common: 5’-GCC TTC TCC TCT AGG CTC GT-3’

Vcre WT: 5’-TAT AGG GCA GAG CTG GAG GA-3’

Vcre Mutant reaction

Vcre Common: 5’-GCC TTC TCC TCT AGG CTC GT-3’

Vcre Mutant: 5’-AGG CAA ATT TTG GTG TAC GG-3’

### Diet and experimental design

Mice were weaned at 21 days of age and placed on a chow diet (Purina Laboratory Rodent Diet 5015). At 8 weeks of age, female LFABP^liv-/-^, LFABP^int-/-^ and LFABP^fl/fl^ control mice were fed a 45% Kcal fat semipurified HFD (D10080402, Research Diets, New Brunswick, NJ) for 12 weeks. The diet composition was described previously [21]. All animal experiments were approved by the Rutgers University Animal Care and Use Committee.

### Body Weight and Body Composition

During the HFD feeding period, the body weight was measured each week. FM and fat-free mass (FFM) were measured using MRI (Echo Medical Systems, LLC., Houston, TX) 1–2 days before starting the feeding protocol, and 1–2 days before sacrificing the mice. The instrument was calibrated each time according to the manufacturer’s instructions. At each time point, 2 measurements were taken for each mouse and averaged.

### Indirect Calorimetry, Activity, and Food Intake

Respiratory exchange ratio (RER), activity, and food intake were assessed using the Oxymax system (Columbus Instruments, Columbus, OH) during weeks 10–11 of the feeding protocol. Mice were placed in an indirect calorimetry chamber (1 mouse per chamber) with food for 48 hours. The first 24 hours were used as an acclimation period, while the second 24-hour period was used for recording RER (VCO2/VO2), activity, and food intake. Energy expenditure (EE) was measured by using the gas exchange measurements as follows: (3.815 + 1.232 × RER) × VO2 [38].

### Intestinal Transit Time

Transit time was measured between weeks 11 and 12 of the HFD feeding period. Prior to the start of the experiment, mice were individually caged. After 2 hours of acclimation, mice were given 250 μL of 6% carmine red and 0.5% methylcellulose (Sigma-Aldrich, St. Louis, MO) in PBS by oral gavage. After gavaging the mice, the cages were then checked every 10 minutes and the time of appearance of the first red fecal pellet was recorded [39,40].

### Total Fecal Excretion

Mice were housed 2–3 per cage. Feces from each cage were collected for 3–4 days between weeks 11 and 12 of the HFD feeding period, dried overnight at 60^○^C, and then weighed. The weight of the feces was converted into Kcal energy excreted and divided by the number of mice in the cage and by the number of days of collection [21]. To control for differences in food intake (energy intake), the averaged energy intake was measured for the mice in the same cage from which the feces were collected. The results of the averaged feces excreted were normalized to their respective averaged 24-hour energy intakes, to generate values of Kcal energy absorbed per mouse per day.

### Treadmill Exercise Protocol

After 12 weeks of HF feeding, exercise endurance was assessed using a treadmill inclined at 25°. One day prior to the test, mice were acclimated by walking at 5 m/min for 5 minutes. For the test, the speed began at 6 m/min for 5 minutes, and then was increased by 3 m/min every 2 minutes. The treadmill had a shock grid at the base, which was kept at a low electrical intensity. When the mice failed to keep up with the treadmill belt, they came in contact with the shock grid. If the mice remained on the shock grid for 5 seconds, they were considered to be exhausted and have fatigue; at this time the mice were removed from the apparatus, and the time to fatigue and total distance traveled were recorded [41,42].

### Oral Glucose Tolerance Tests (OGTT)

During week 11 of the HF feeding protocol, mice were fasted for 6 hours prior to the OGTT experiments. Blood was drawn from the tail vein, and baseline glucose was measured using an Accu-Check monitor. An oral gavage of 2 g glucose/Kg body weight was then administered, and blood was sampled at time points of 30, 60, 90, and 120 minutes.

### Tissue Preparation

At the end of the HF feeding period, mice were fasted for 16 hours prior to sacrifice. Before sample collection mice were anesthetized with ketamine-xylazine-acepromazine (80, 100, 150 mg/kg intraperitoneally [IP], respectively). Epididymal and inguinal fat pads, and livers were removed, weighed, immediately placed on dry ice, and stored at −80°C for further analysis. The SI from stomach to cecum was removed, measured lengthwise, rinsed with 60 mL of ice-cold 0.1M NaCl, and opened longitudinally. Intestinal mucosa was scraped with a glass microscope slide into tared tubes on dry ice to be further used for mRNA extraction, protein extraction, or lipid extraction.

### Blood Preparation and Analysis

At time of sacrifice, whole blood was used to measure glucose (Accu-Check, Roche Diagnostics). Plasma was isolated after centrifugation for 6 minutes at 4000 rpm, and stored at −80°C for further analysis. ELISA kits were used to measure plasma insulin (Millipore), leptin (Millipore), and adiponectin (Sigma-Aldrich). Plasma cholesterol and free fatty acid (FFA) were measured colorimetrically using Cell Biolabs kits, and TG was measured colorimetrically using a Cayman kit. Adiponectin and leptin indices were calculated by dividing adiponectin or leptin levels by the total FM determined by MRI.

### RNA Extraction and Real-Time PCR

Total mRNA was extracted from SI mucosa and liver and analyzed as previously described [20,21]. Primer sequences (**Supplemental Table 1**) were obtained from Primer Bank (Harvard Medical School QPCR Primer Database). The efficiency of PCR amplifications was checked for all primers to confirm similar amplification efficiency. Real time PCR reactions were performed in triplicate using an Applied Biosystems StepOne Plus instrument. Each reaction contained a suitable amount of cDNA, 250nM of each primer, and 12.5 μL of SYBR Green Master Mix (Applied Biosystems, Foster City, CA) in a total volume of 25 μL. Relative quantification of mRNA expression was calculated using the comparative Ct method, normalized to endogenous TATA-binding protein.

### Lipid Extraction and Metabolites Analysis

Mucosa and liver samples collected as described above were used for lipid extraction and thin layer chromatography analysis, as described previously [21,27,43,44].

### VLDL-TG Secretion Measurement

After 12 weeks of HF feeding, mice were fasted for 6 hrs. Then an IP injection of Tyloxapol (500 mg/kg body weight) was administered to block lipolysis of TG via inhibition of lipoprotein lipase. At time 60, 90, 120, 150, and 180 minutes after injection, 15 μl of blood was collected from conscious mice via the tail vein. Blood TG levels were measured using a Cardiochek instrument (Polymer Technology Systems, Inc. Zionsville, IN).

### Oral Fat Tolerance Test (OFTT)

OFTT was performed as described previously [21]. After 12 weeks of HF feeding, mice were fasted for 6 hours. Time 0 blood was taken from conscious mice via the tail vein and then an IP injection of Tyloxapol (500 mg/kg body weight) was administered. After 30 min, an orogastric gavage of 300 μL of olive oil was given. Blood was taken at time 1, 2, 3, and 4 hours. Blood TG levels were measured using 15 μl of whole blood from the tail vein using a Cardiochek instrument (Polymer Technology Systems, Inc. Zionsville, IN).

### FA Oxidation Measurements

FA oxidation rates in liver homogenates were measured as detailed by Huynh and colleagues [45]. Briefly, upon sacrifice, livers (approximately 200 mg) were gently homogenized with a Potter–Elvehjem homogenizer for 5 strokes on ice, using 5× the weight of the samples (wet weight) of sucrose–Tris–EDTA, and the homogenates were centrifuged at 420 × g for 10 min at 4°C; the supernatants were then incubated for 30 min at 37°C with 370 µl of reaction mixture containing 1.6 µCi of ^14^C oleate solubilized in 0.7% bovine serum albumin (BSA), and 500 µM palmitate. ^14^CO_2_ generated from the reaction was released by adding 200 µl of 1M perchloric acid and absorbed onto a piece of filter paper in the tube cap soaked with 10 µl of 1M NaOH. The filter paper and ^14^C-labeled acid soluble metabolites (ASMs) in the reaction mixture were assessed for radioactivity by scintillation counting.

### Tissue FA Uptake Assay

FA uptake into different tissues was measured as described by others [46–48] with minor modifications. Overnight-fasted mice received an orogastric gavage of ^14^C Oleic acid (2.5 μCi) in 200 μL olive oil. Mice were anesthetized with ketamine-xylazine-acepromazine (80, 100, 150 mg/kg IP, respectively) 4 hours after the oral lipid load. Blood samples were drawn from anesthetized mice, and plasma was extracted by adding 9% of perchloric acid and then centrifuged for 1 min at 16,000 × g. Liver, gastrocnemius muscle, and epidedmal and inguinal white adipose tissue were removed, rinsed with NaCl and blotted dry. Tissues were weighed and EcoLume cocktail Counting Scintillant was added. Total radioactivity was measured using scintillation counting. SI were also excised, washed with 10 mL 0.8% NaCl and divided into 2 parts, the proximal intestine (PI) and distal intestine (DI). EcoLume cocktail Counting Scintillant was added. Both the intestinal tissues and the non-absorbed luminal content (in NaCl) were examined for ^14^C activity to determine the amount of absorbed versus non-absorbed FA present in the intestinal tract.

### Statistical Analysis

The results were analyzed using GraphPad Prism 10 version 2. Data are expressed as mean ± SD. Statistical comparisons were made by a 2-sided Student’s t-test versus WT. Differences were considered significant at *P* <0.05 (symbols * and # <0.05; ** and ## <0.01; and *** and ### <0.001).

## 3. Results

### Ablation of LFABP was specific to the liver in the LFABP^liv-/-^ mice and specific to the intestine in the LFABP^int-/-^ mice

Tissue-specific ablation of LFABP was confirmed by Western blot, with LFABP^Liv-/-^ mice expressing LFABP only in the intestine (**Figure 1A**) and LFABP^int-/-^ mice expressing LFABP only in the liver (**Figure 1B**). Control LFABP^fl/fl^ mice expressed LFABP in both the liver and the intestine as expected (**Figure 1**).

**Figure 1.**
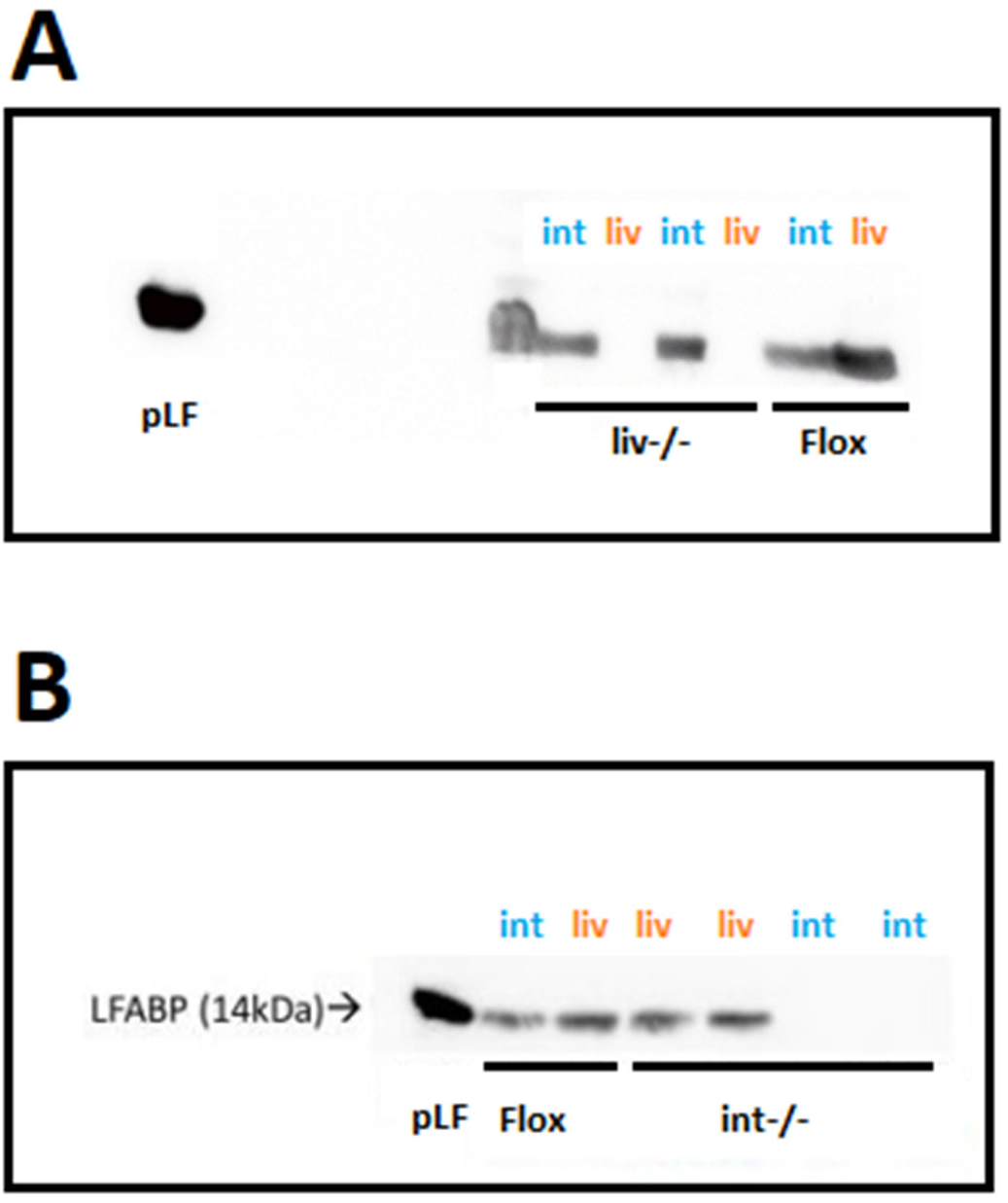
Confirmation of the expression of LFABP in the liver and intestine of LFABP^fl/fl^, LFABP^liv-/-^, and LFABP^int-/-^ mice. A, Western blot analysis confirms the ablation of liver-LFABP in LFABP^liv-/-^ (liv-/-) mice; B, Western blot analysis confirms the ablation of intestine-LFABP in LFABP^int-/-^ (int-/-) mice. int, intestine; LFABP, liver fatty acid-binding protein; LFABP^fl/fl^, floxed liver fatty acid-binding protein; LFABP^int-/-,^ intestine-specific liver fatty acid-binding protein knockout; LFABP^liv-/-^, liver-specific liver fatty acid-binding protein knockout; liv, liver; pLF, purified liver fatty acid-binding protein.

### cKO LFABP mice have greater body weight and FM compared with the control floxed mice

At 2 months of age, LFABP^fl/fl^, LFABP^liv-/-^, and LFABP^int-/-^ mice were challenged with a 45% Kcal fat HFD. After 12 weeks of HF feeding, the body weights (**Figure 2A**) and body weight gain (**Figure 2B**) of both groups of cKO mice were significantly higher than those of their LFABP^fl/fl^ counterparts. LFABP^liv-/-^ mice and LFABP^int-/-^ mice also had a greater FM % than LFABP^fl/fl^ mice (**Figure 2C**). The body weight of LFABP^int-/-^ mice was significantly higher than LFABP^fl/fl^ control mice starting at week 4 on the HFD; while for LFABP^liv-/-^ mice body weight was significantly higher starting at week 8 on the HFD.

**Figure 2.**
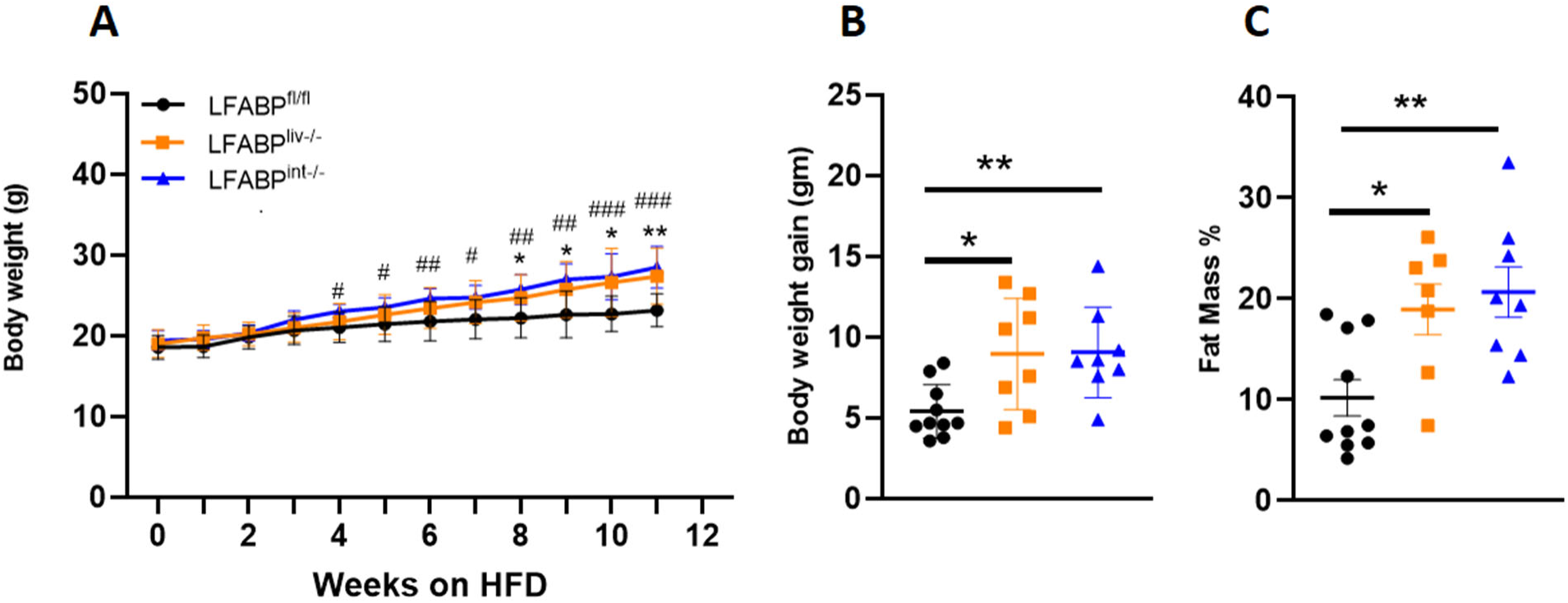
Body weight, body weight gain, and fat mass % for LFABP^fl/fl^ (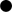), LFABP^liv-/-^ (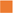) and LFABP^int-/-^ (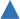) mice after 12 weeks of 45% Kcal HF feeding. A, Body weights (n = 8–10); B, Body weight gain (n = 8–10); C, Fat mass percentage (n = 8–9). Data are given as mean ± SD, analyzed using Student’s t-test. *, *P* < 0.05 and **, *P* < 0.01 for LFABP^liv-/-^ versus LFABP^fl/fl^; #, *P* < 0.05, ##, *P* < 0.01 and ###, *P* < 0.001 for LFABP^int-/-^ versus LFABP^fl/fl^ in figure A. *, *P* < 0.05 and **, *P* < 0.01 for LFABP cKO mice versus LFABP^fl/fl^ in figures B and C. LFABP, liver fatty acid-binding protein; LFABP^fl/fl^, floxed liver fatty acid-binding protein; LFABP^int-/-^, intestine-specific liver fatty acid-binding protein knockout; LFABP^liv-/-^, liver-specific liver fatty acid-binding protein knockout.

### The ablation of LFABP from either the liver or intestine does not alter net energy absorption, intestinal transit times, or EE

Mice were placed into the Oxymax system for indirect calorimetry measurements and to assess food intake. Feces were collected to measure fecal mass excreted over 24 hours. Despite the observed differences in body weight and body composition between LFABP^fl/fl^ and both LFABP^liv-/-^ and LFABP^int-/-^ female mice, there were no significant alterations in the calories consumed and energy absorbed (**Figure 3A**). Additionally, there were no differences in the intestinal transit times between the LFABP^liv-/-^ mice and the LFABP^fl/fl^ control mice (**Figure 3B**); a trend toward slower transit time was seen for the LFABP^int-/-^ mice but this did not reach statistical significance (**Figure 3B**). There were also no differences between groups in either 24-hour RER or EE (**Figure 3C and D**).

**Figure 3.**
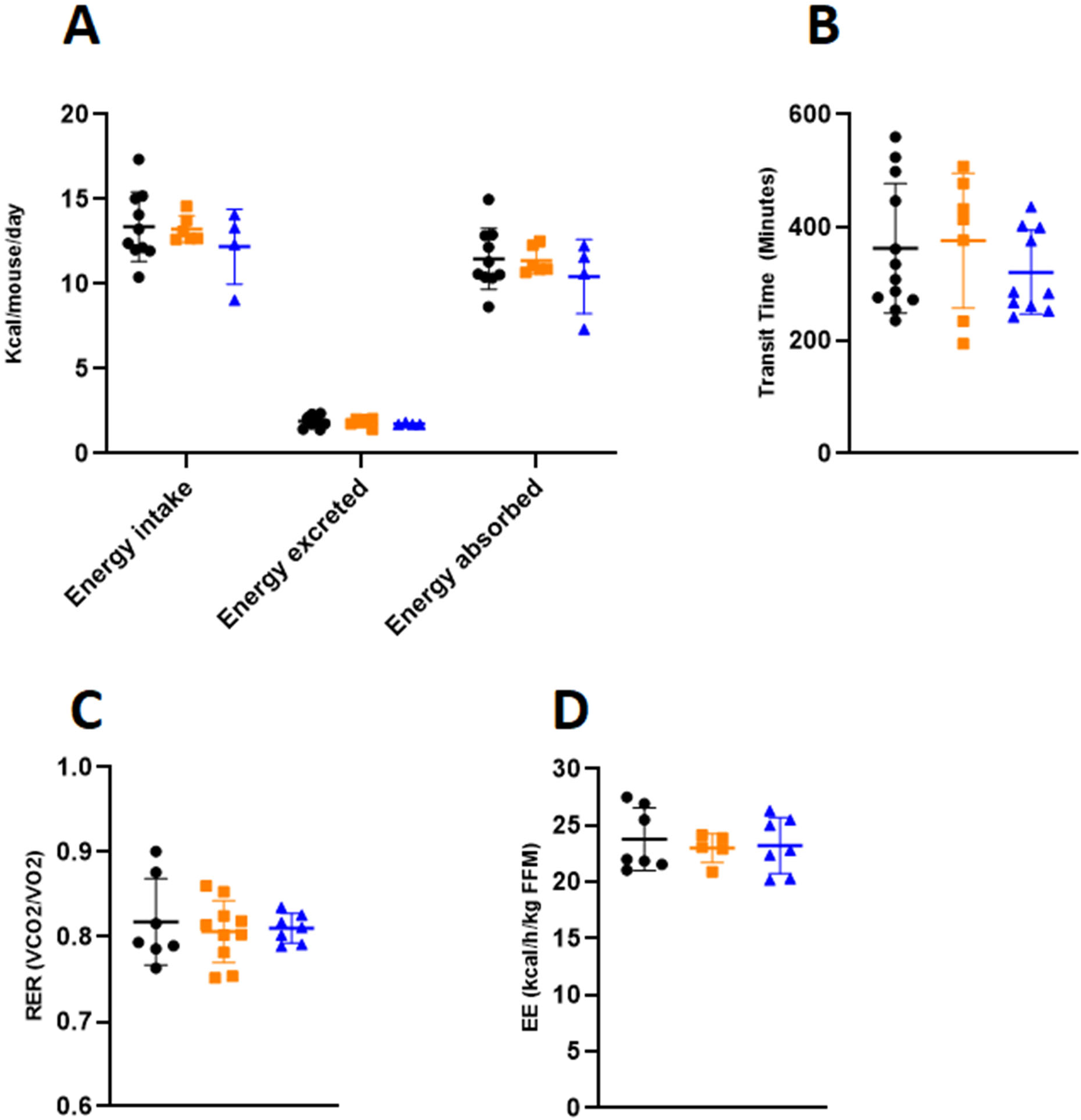
Food intake, intestinal transit times, RER and EE in LFABP^fl/fl^ (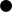), LFABP^liv-/-^ (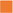) and LFABP^int-/-^ (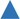) mice after 12 weeks of 45% Kcal HF feeding. A, 24-hour energy intake, feces excreted and energy absorbed (n = 4–10); B, intestinal transit time (n = 7–12); C, 24-hour RER (n = 7–10); B, EE (n = 5–7). Data are given as mean ± SD, analyzed using Student’s t-test. EE, Energy expenditure; LFABP^fl/fl^, floxed liver fatty acid-binding protein; LFABP^int-/-^, intestine-specific liver fatty acid-binding protein knockout; LFABP^liv-/-^, liver-specific liver fatty acid-binding protein knockout; RER, Respiratory exchange ratio.

### LFABP^liv-/-^ and LFABP^int-/-^ mice retain their exercise capacity upon HFD feeding relative to LFABP^fl/fl^ control mice

Both spontaneous and induced physical activity parameters were assessed in the intestine-specific and liver-specific LFABP cKO mice. No alterations in 24-hour spontaneous activity (**Figure 4A**) were noted in either cKO mouse group. However, both LFABP^liv-/-^ and LFABP^int-/-^ mice displayed higher exercise endurance capacity relative to their LFABP^fl/fl^ controls (**Figure 4B**).

**Figure 4.**
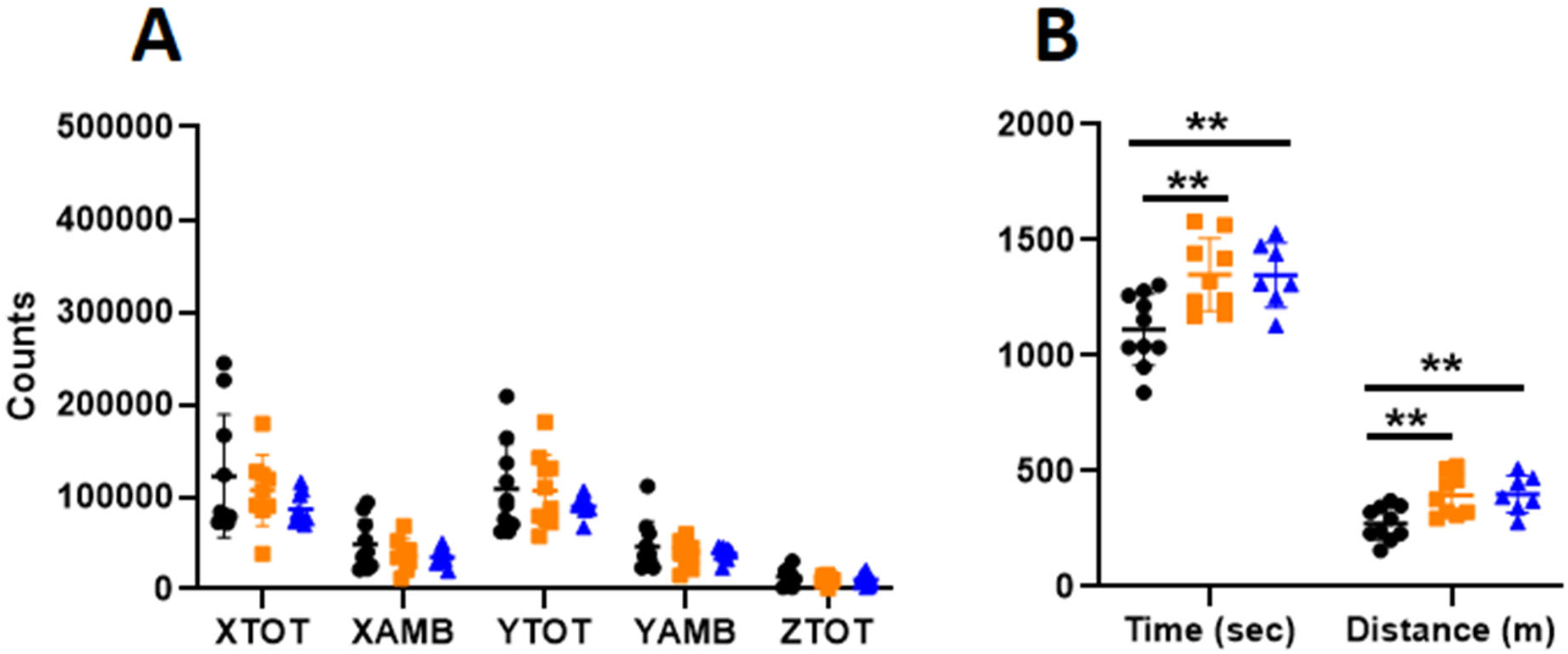
Analyses of spontaneous activity and exercise endurance capacity for LFABP^fl/fl^ (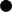), LFABP^liv-/-^ (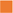) and LFABP^int-/-^ (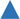) mice after 12 weeks of 45% Kcal HF feeding. A, 24-hour spontaneous activity (n = 8–10); B, exercise endurance running time and distance (n = 7–10). Data are given as mean ± SD, analyzed using Student’s t-test. **, *P* < 0.01 for LFABP cKO mice versus LFABP^fl/fl^ mice. AMB, ambulatory; LFABP^fl/fl^, floxed liver fatty acid-binding protein; LFABP^int-/-^, intestine-specific liver fatty acid-binding protein knockout; LFABP^liv-/-^, liver-specific liver fatty acid-binding protein knockout; TOT, total.

### Liver- and intestine-LFABP cKO mice do not display alterations in plasma markers of energy balance

Despite increased adiposity, LFABP^liv-/-^ and LFABP^int-/-^ mice showed no differences in blood glucose concentrations at any time point after the gavage and no differences in fasting plasma insulin, when compared with LFABP^fl/fl^ mice (**Figure 5 A and B**, **Table 1**). While fasting plasma leptin was higher in LFABP^liv-/-^ mice relative to LFABP^fl/fl^ mice the leptin index, which factors in FM, revealed no significant difference between these 2 groups. Fasting plasma leptin showed a nonsignificant increase in LFABP^int-/-^ mice however as with the LFABP^liv-/-^, the leptin index was comparable to the control mice (**Table 1**). Adiponectin level and index showed no difference between LFABP^liv-/-^ mice and their control mice; there were also no significant changes in the plasma levels of non-esterified fatty acids (NEFA), TG, and cholesterol in female LFABP^liv-/-^ mice when compared with the floxed WT control mice.

**Figure 5.**
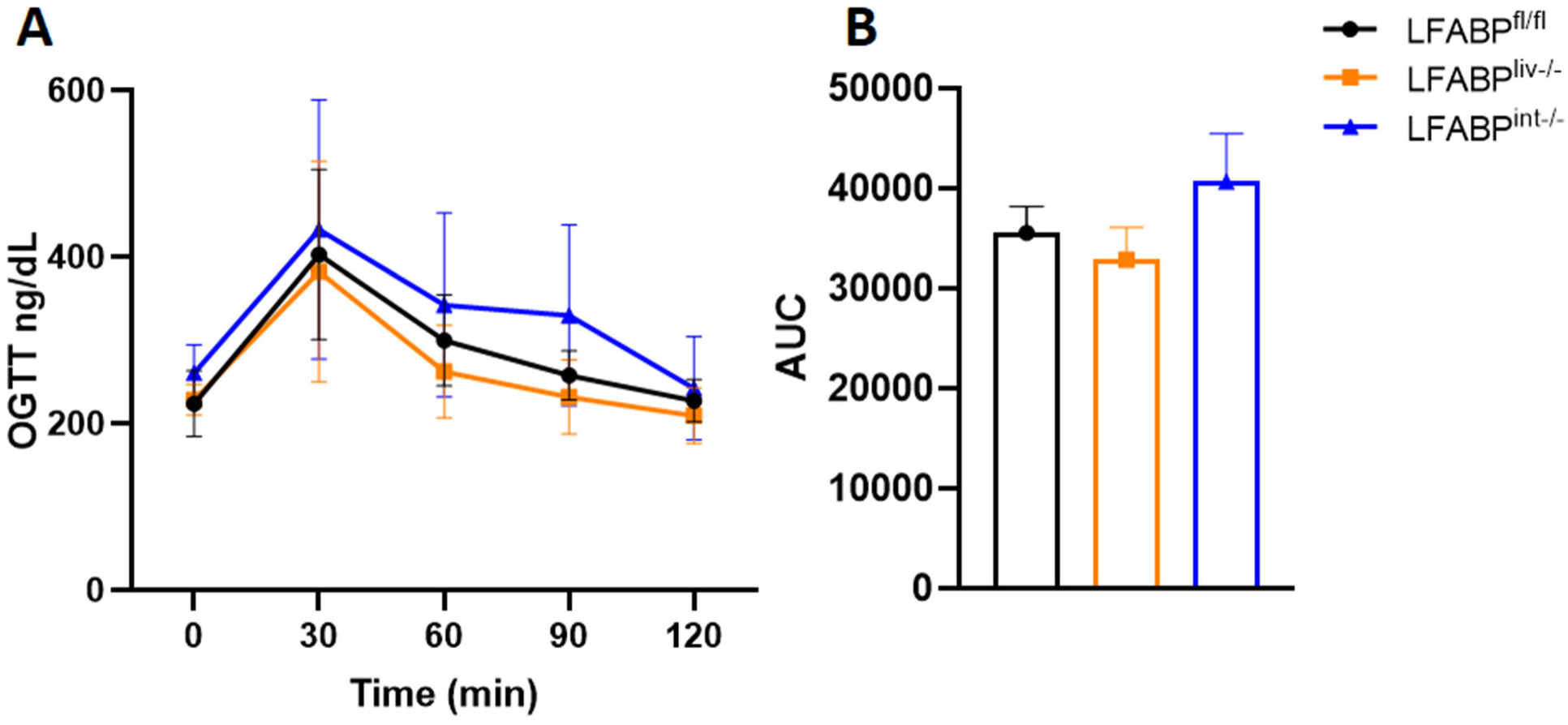
OGTT for fasted LFABP^fl/fl^ (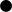), LFABP^liv-/-^ (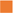) and LFABP^int-/-^ (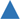) mice after 12 weeks of 45% Kcal HF feeding. A, OGTT (n = 7–13); B, OGTT area under the curve (AUC) (n = 7-13) Data are given as mean ± SD for figure A and mean ± SE for figure B, analyzed using Student’s t-test. LFABP, liver fatty acid-binding protein; LFABP^fl/fl^, floxed liver fatty acid-binding protein; LFABP^int-/-^, intestine-specific liver fatty acid-binding protein knockout; LFABP^liv-/-^, liver-specific liver fatty acid-binding protein knockout; OGTT, Oral glucose tolerance test.

**Table 1.** Plasma analyses for LFABP^fl/fl^, LFABP^liv-/-^ and LFABP^int-/-^ mice after 12 weeks of 45% Kcal HF feeding

|  | LFABP<sup>fl/fl</sup> | LFABP<sup>liv-/-</sup> | LFABP<sup>int-/-</sup> |
| --- | --- | --- | --- |
| Glucose (ng/dL) | 240.1 ± 24.68 | 228.22 ± 18.34 | 260.86 ± 33.84 |
| Insulin (ng/mL) | 0.37 ± 0.09 | 0.40 ± 0.04 | 0.30 ± 0.05 |
| Leptin (ng/mL) | 4.74 ± 2.34 | 16.81 ± 8.20** | 6.87 ± 4.41 |
| Leptin index | 1.54 ± 0.60 | 2.05 ± 0.22 | 1.03 ± 0.24 |
| Adiponectin (ng/mL) | 14406.04 ± 5931.61 | 14946.52 ± 2494.73 | ND |
| Adiponectin index | 4322.25 ± 2787.70 | 2246.00 ± 1123.28 | ND |
| NEFA (μM) | 357.74 ± 153.58 | 402.42 ± 210.23 | ND |
| TG (mg/dL) | 19.34 ± 11.44 | 16.70 ± 11.31 | ND |
| Cholestrol (μM) | 1298.41 ± 468.77 | 1414.29 ± 349.85 | ND |

Data are given as mean ± SD, analyzed using Student's t-test. \*\*, $P < 0.001$ for LFABP<sup>liv-/-</sup> mice versus LFABP<sup>fl/fl</sup> mice. n = 6–11 for all groups.
HF, high-fat; LFABP, liver fatty acid-binding protein; LFABP<sup>fl/fl</sup>, floxed liver fatty acid-binding protein; LFABP<sup>int-/-</sup>, intestine-specific liver fatty acid-binding protein knockout; LFABP<sup>liv-/-</sup>, liver-specific liver fatty acid-binding protein knockout; ND, not determined; NEFA, non-esterified fatty acids.

### Hepatic lipid handling in LFABP cKO mice

Since mice challenged with chronic HF feeding progressively develop fatty liver in addition to obesity, we investigated the effects of long-term HFD combined with liver- or intestinal-LFABP ablation on the liver phenotype. While absolute liver weight was significantly higher in LFABP^liv-/-^ mice, the liver weight/body weight ratio was significantly somewhat lower relative to LFABP^fl/fl^ control mice. The liver weight of LFABP^int-/-^ mice was also greater but the liver weight/ body weight ratio was comparable to the control mice (**Figure 6A and B**). Despite the greater obesity of the cKO mice, hepatic TG levels in overnight fasted LFABP^liv-/-^ were comparable to their LFABP^fl/fl^ control counterparts; no differences were found in other lipid species as well, including CE, FFA, PL, cholesterol, diglycerides (DG), and monoglycerides (MG), when compared with the LFABP^fl/fl^ controls (**Figure 6C and D**). No significant differences were found in the rate of hepatic FA oxidation, assessed by quantifying ^14^CO_2_ and ^14^C-labeled ASMs in liver homogenates, for either cKO mice when compared with the floxed control mice (**Figure 6E and F**).

**Figure 6.**
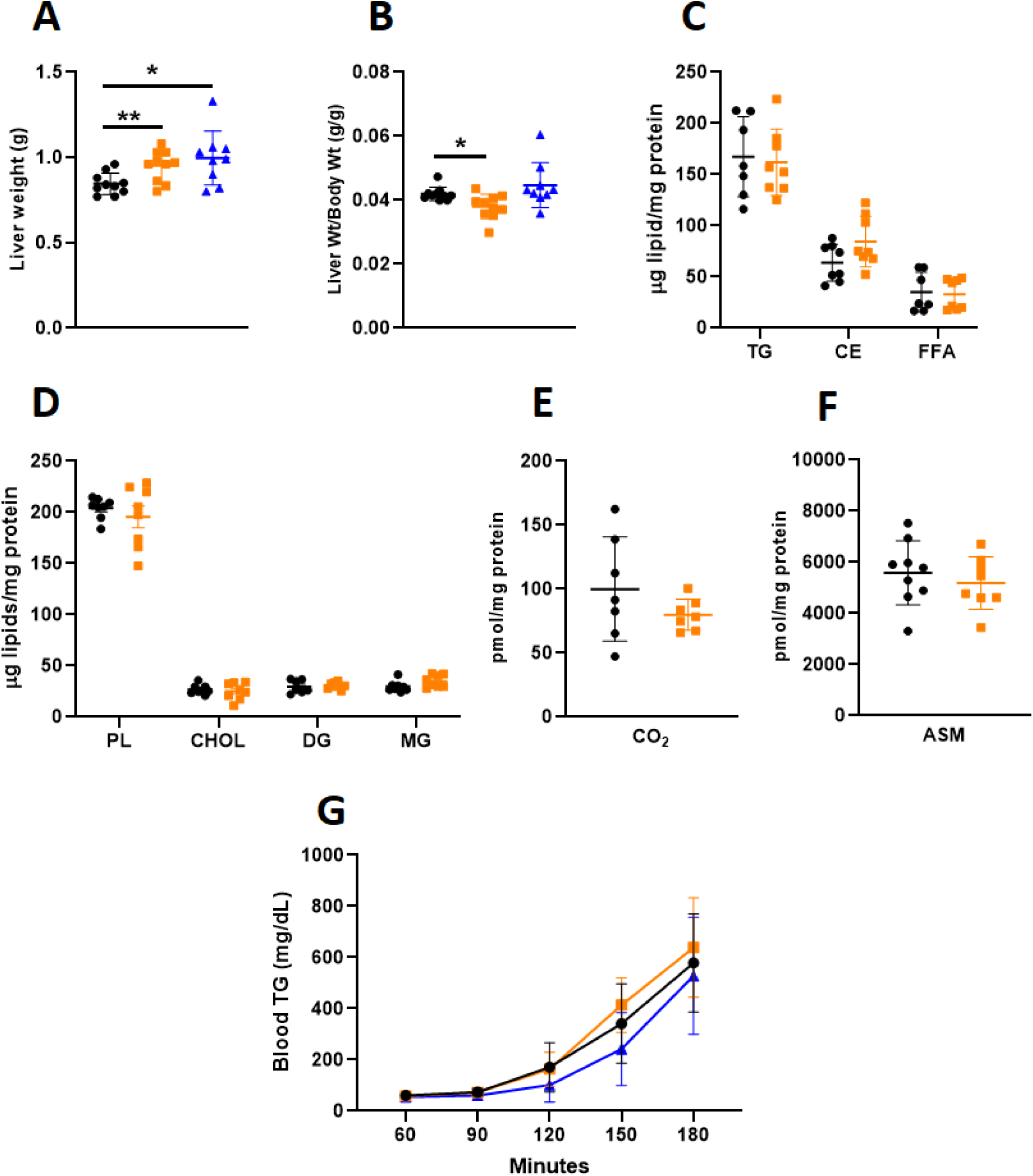
Liver weights and hepatic lipid handling in LFABP^fl/fl^ (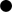), LFABP^liv-/-^ (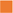) and LFABP^int-/-^ (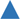) mice after 12 weeks of 45% Kcal HF feeding. A, Liver weight (n = 9–10); B, liver weight/body weight ratio (n = 9–10); C, hepatic neutral lipids (TG, CE and FFA) levels (n = 7–8); D, hepatic lipid species (n = 6–8); E, FA oxidation rate, ^14^CO_2_ production (n = 8–9); F, FA oxidation rate, ^14^C-labeled ASMs (n = 10–11); G, blood VLDL-TG level (n = 5–14); Data are given as mean ± SD, analyzed using Student’s t-test. *, *P* < 0.05 and **, *P* < 0.01 for cKO LFABP versus LFABP^fl/fl^. ASM, Acid soluble metabolite; CHOL, cholesterol; CE, cholesteryl ester; DG, diglyceride; FA, fatty acid; LFABP^fl/fl^, floxed liver fatty acid-binding protein; LFABP^int-/-^, intestine-specific liver fatty acid-binding protein knockout; LFABP^liv-/-^, liver-specific liver fatty acid-binding protein knockout; MG, monoglyceride; PL, phospholipid; TG, triglyceride.

To investigate the mechanisms underlying the lower-than-expected hepatic lipid accumulation in the obese LFABP^liv-/-^ mice, plasma levels of TG-rich VLDL were assessed as described under Methods. The LFABP^liv-/-^ mice showed a trend toward higher VLDL-TG secretion, while LFABP^int-/-^ mice showed a trend toward lower VLDL-TG secretion, although neither difference reached statistical significance (**Figure 6G**).

### Intestinal lipid handling in LFABP cKO mice

We assessed effects of liver-specific or intestine-specific ablation of LFABP on the intestine. The ratio of intestinal length to body weight was not different for LFABP^liv-/-^ mice but was lower for the LFABP^int-/-^ relative to their controls (**Figure 7A**). The intestine of LFABP^liv-/-^ mice displayed a significantly higher accumulation of TG and reduction of PL relative to LFABP^fl/fl^ (**Figure 7B**). Since the redistribution of intestinal lipids could result from changes in TG-rich chylomicron secretion, OFTTs were performed to assess whether the secretion of TG-rich chylomicrons was affected by the ablation of LFABP specifically from the liver or the intestine. The intestine-specific ablation of LFABP resulted in a substantial `∼`50% decline in chylomicron secretion (**Figure 7C, D**), and the LFABP^liv-/-^ mice showed a smaller, 19% reduction in chylomicron secretion (**Figure 7C, D**).

**Figure 7.**
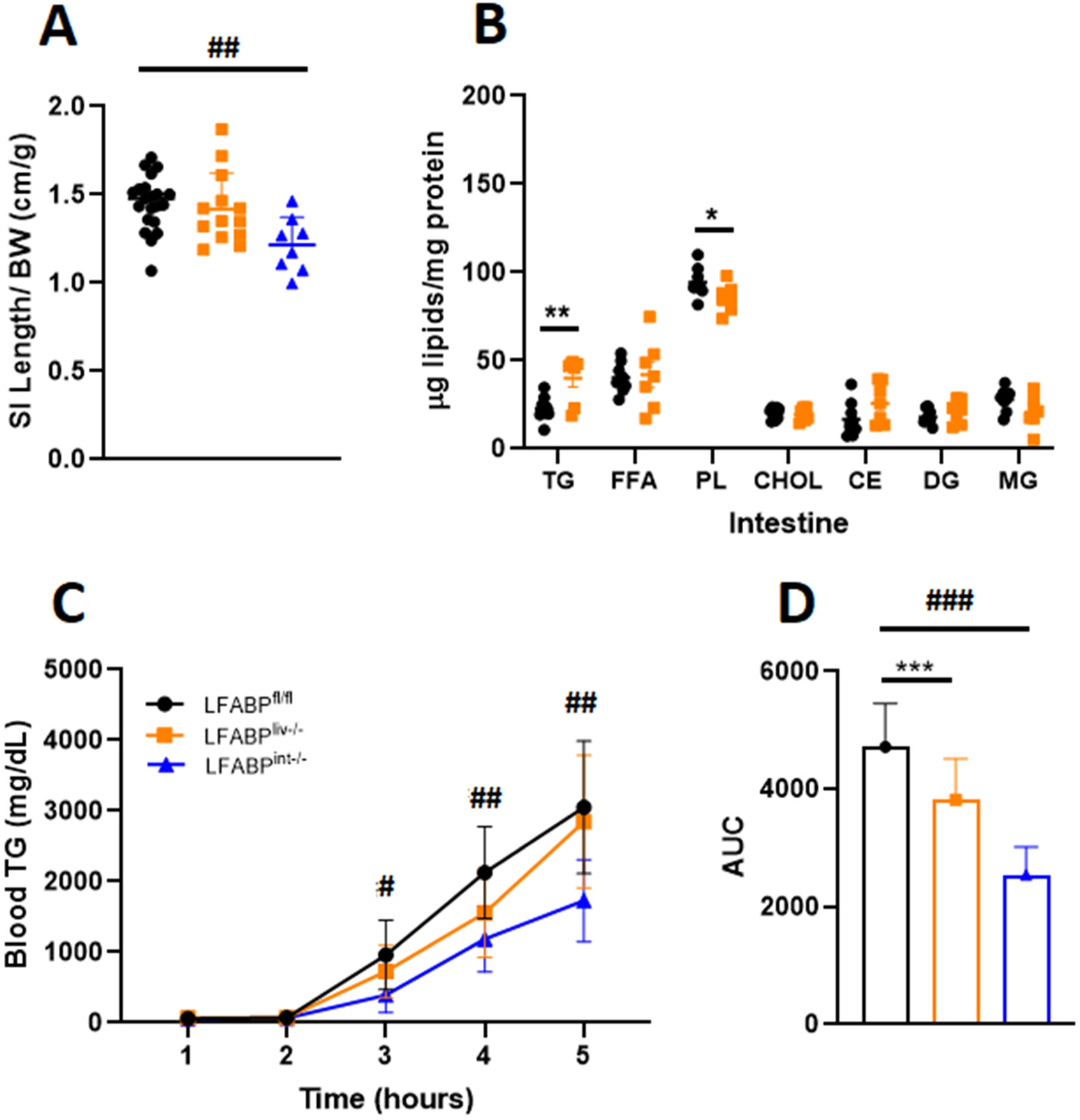
Intestinal lipid handling in LFABP^fl/fl^ (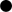), LFABP^liv-/-^ (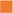) and LFABP^int-/-^ (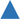) mice after 12 weeks of 45% Kcal HF feeding. A, Intestine length/BW ratio (n = 8-21); B, intestinal lipid species concentrations (n = 6–9); C, intestinal chylomicron secretion rates (blood TG levels) (n = 6–8); D, intestinal chylomicron secretion rates AUC (n = 6-8). Data are given as mean ± SD, analyzed using Student’s t-test. *, *P* < 0.05; **, *P* < 0.01 and ***, *P* < 0.001 for LFABP^liv-/-^ versus LFABP^fl/fl^. #, *P* < 0.05; ##, *P* < 0.01 and ###, P < 0.001 for LFABP^int-/-^ versus LFABP^fl/fl^. AUC, area under the curve; BW, body weight; CHOL, cholesterol; CE, cholesteryl ester; DG, diglyceride; FFA, free fatty acid; LFABP^fl/fl^, floxed liver fatty acid-binding protein; LFABP^int-/-^, intestine-specific liver fatty acid-binding protein knockout; LFABP^liv-/-^, liver-specific liver fatty acid-binding protein knockout; MG, monoglyceride; PL, phospholipid; TG, triglyceride; SI, Small intestine.

### Tissue uptake of oral FA is modulated by LFABP conditional ablation

In LFABP^liv-/-^ mice, tissue FA uptake was significantly reduced in the liver and proximal intestine, and displayed a trend toward higher FA uptake in adipose tissue relative to the LFABP^fl/fl^ control mice (**Figure 8**). The radioactivity in other tissues, feces, and blood did not differ from control mice. Intestine-specific LFABP ablation resulted in a significant reduction in intestinal FA uptake when compared with their floxed control mice. There was also a reduction in the level of labeled FA in the blood of LFABP^int-/-^ mice relative to LFABP^fl/fl^ mice (**Figure 8**).

**Figure 8.**
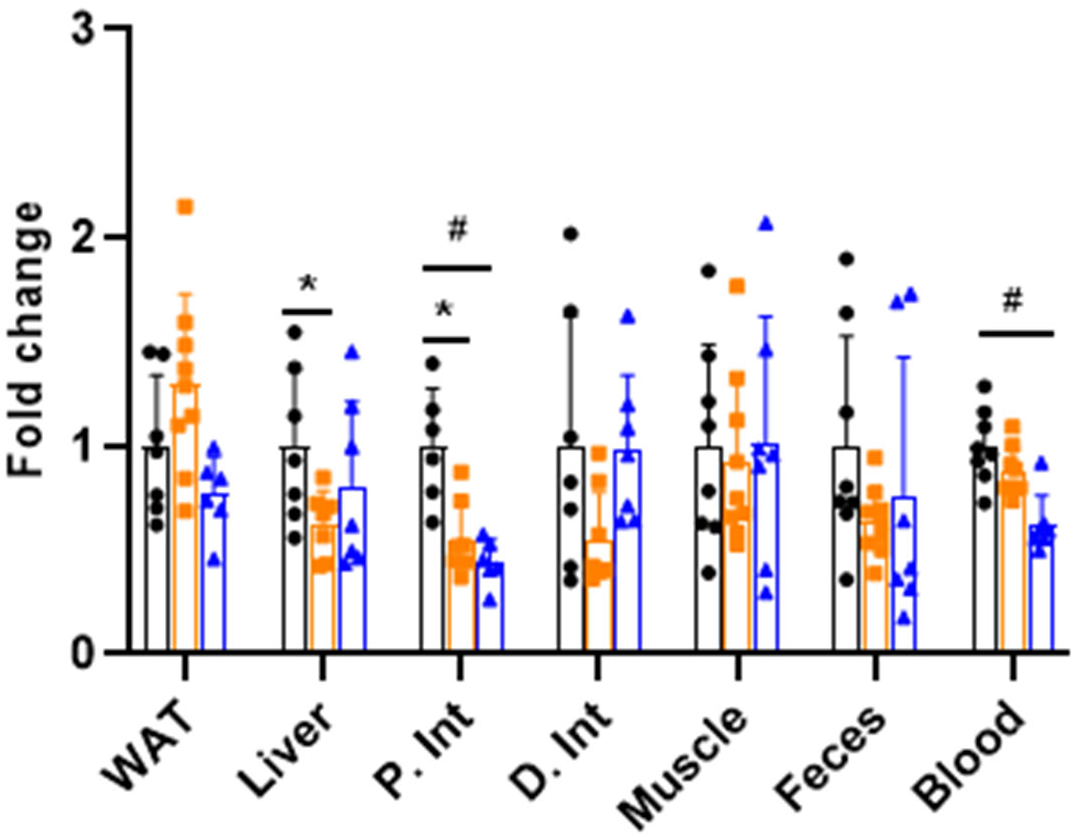
Tissue FA uptake after oral administration of ^14^C-oleic acid to 12-week HFD-fed LFABP^fl/fl^ (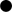), LFABP^liv-/-^ (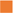) and LFABP^int-/-^ (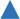) mice following an overnight fast. FA uptake into WAT, liver, P. Int, D. Int, gastrocnemius muscle, ^14^C-oleic acid appearance in the feces and the blood (n = 6–7). Data are given as mean ± SD, analyzed using Student’s t-test. *, *P* < 0.05 for LFABP^liv-/-^ versus LFABP^fl/fl^. #, *P* < 0.05 for LFABP^int-/-^ versus LFABP^fl/fl^. D. Int, distal intestine; LFABP^fl/fl^, floxed liver fatty acid-binding protein; LFABP^int-/-^, intestine-specific liver fatty acid-binding protein knockout; LFABP^liv-/-^, liver-specific liver fatty acid-binding protein knockout; P. Int, proximal intestine; WAT, white adipose tissue.

### The expression of lipid metabolism genes in liver and intestine of LFABP cKO mice

Intestine-specific ablation of LFABP led to 50% or greater reductions in the hepatic expression of several genes involved in lipid and fatty acid transport, including *Fatp2* and *Scp2* (**Figure 9A**). Genes involved in lipid and fatty acid synthesis as well as fatty acid oxidation, also showed lower expression levels in the LFABP^int-/-^ liver, including *Fasn*, *Scd1*, *Cpt1a*, *Cpt2*, and *Acox1* (**Figure 9B**). Effects of liver-specific LFABP knockout had little effect on the expression of liver lipid-related genes; while a few genes were significantly modulated, none of these effects reached a 2-fold level of change (**Figure 9A, B**). In LFABP^liv-/-^ mice no changes in the hepatic expression of the transcriptional regulators *Ppar-α*, *Hnf1α,* and *Hnf4α* or the signaling molecule fibroblast growth factor (*Fgf21*) were found, while in LFABP^int-/-^ a significant reduction in *Ppar-α* expression was noted (**Figure 9C**). Little or no change was observed in genes involved in intestinal lipid metabolism in the LFABP^liv-/-^ mouse (**Figure 9D**).

**Figure 9.**
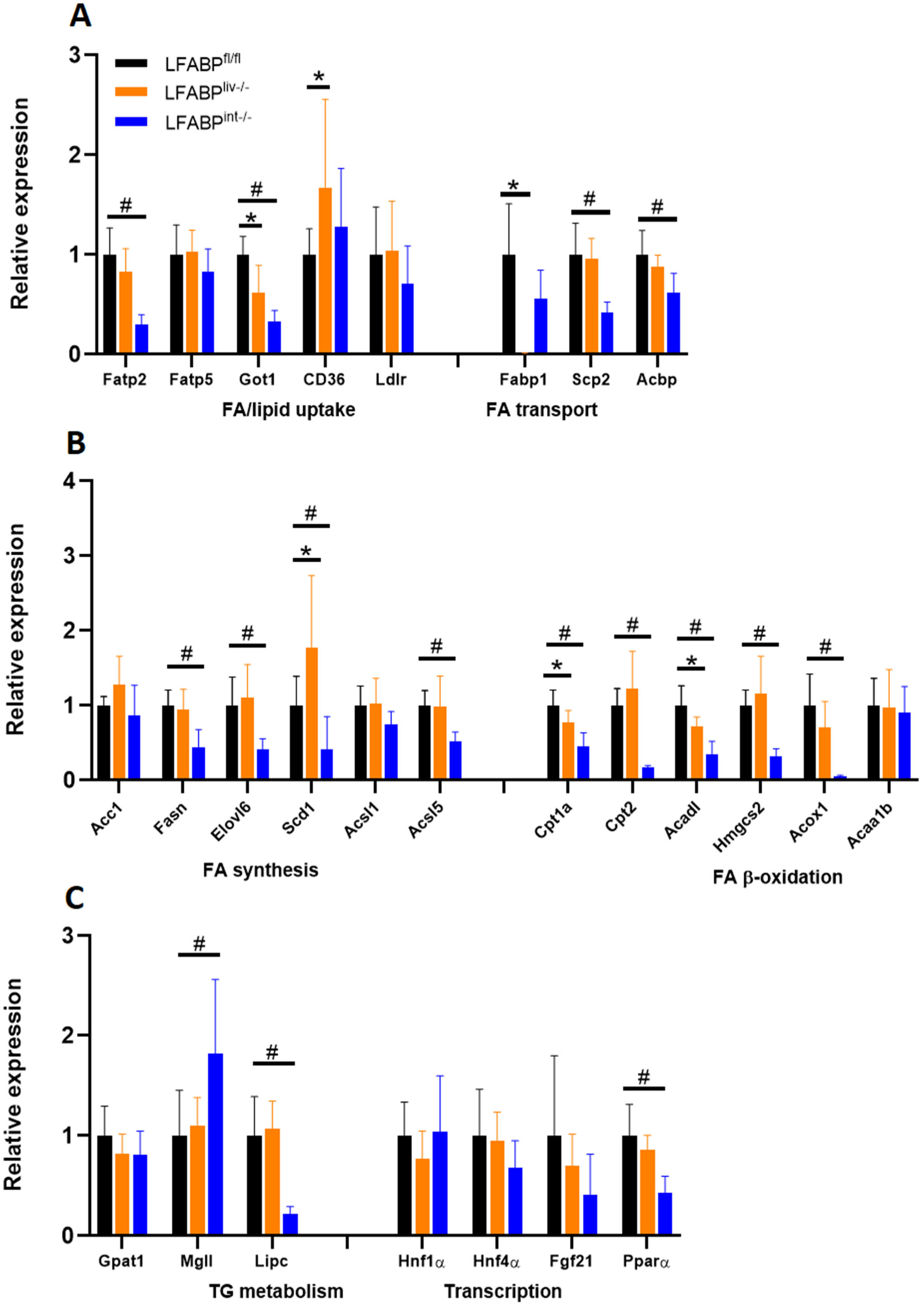
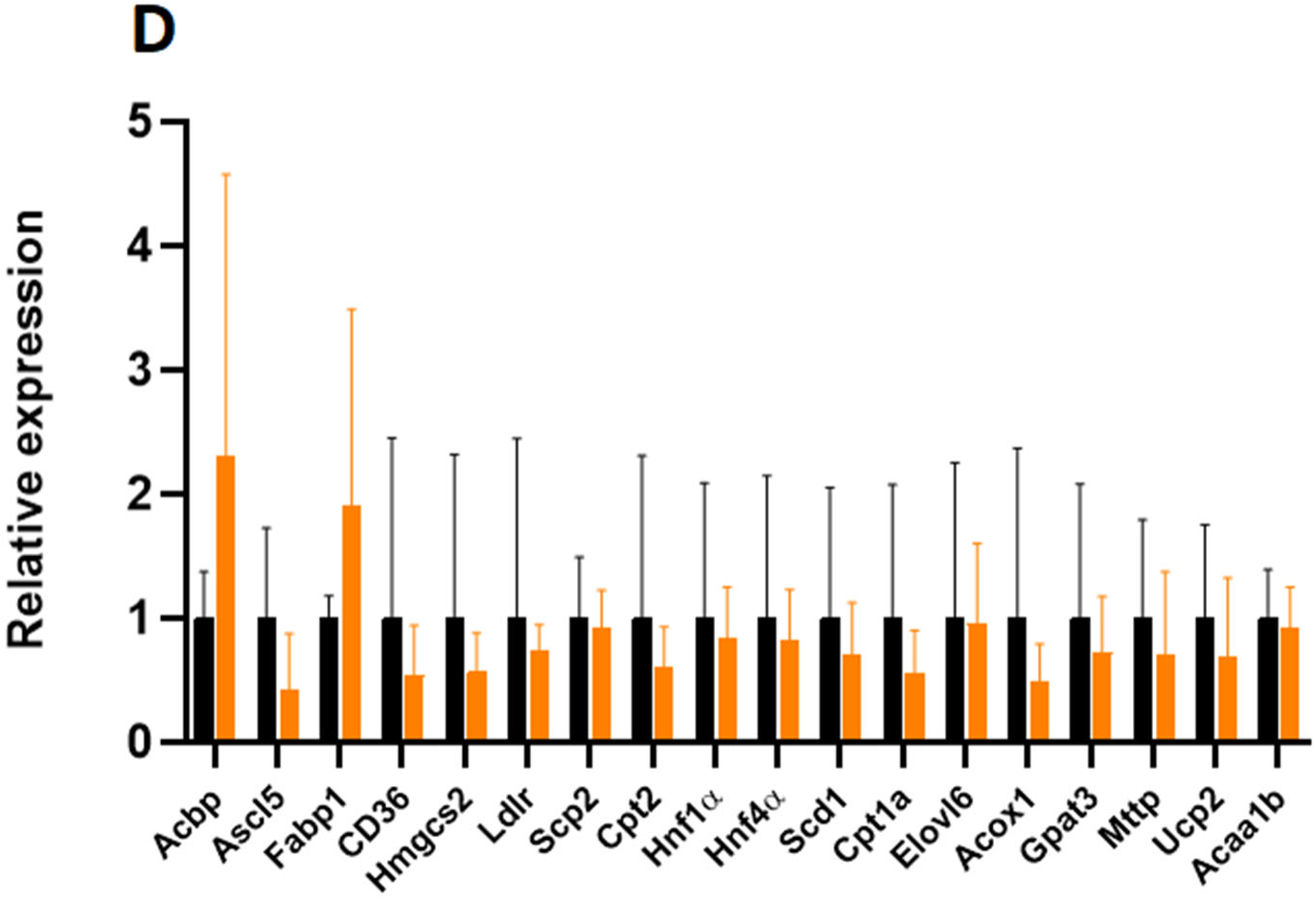
Relative quantitation of mRNA expression of genes involved in liver and intestinal lipid metabolism in 45% Kcal fat HF fed LFABP^fl/fl^ and cKO mice. A, Expression of genes involved in hepatic FA/lipid uptake and FA transport (n = 4–9); B, Expression of genes involved in hepatic FA synthesis and oxidation (n = 4–9); C, Expression of genes involved in hepatic TG metabolism and expression of transcriptional genes (n = 4–9); D, expression of genes involved in intestinal lipid metabolic pathways (n = 5–6); Data are given as mean ± SD, analyzed using Student’s t-test. *, *P* < 0.05 for LFABP^liv-/-^ versus LFABP^fl/fl^, #, *P* < 0.05 for LFABP^int-/-^ versus LFABP^fl/fl^. LFABP^fl/fl^, floxed liver fatty acid-binding protein; LFABP^int-/-^, intestine-specific liver fatty acid-binding protein knockout; LFABP^liv-/-^, liver-specific liver fatty acid-binding protein knockout.

## 4. Discussion

We reported that male LFABP^-/-^ mice become heavier and fatter than WT mice when challenged with chronic HFD feeding [21,23,27,36]; similar increases in body weight and fat mass were found in female mice as well, in agreement with observations in a separate line of LFABP^-/-^ mice [35,49]. The higher body weight gain in the LFABP^-/-^ mice was partly due to increased caloric intake and feeding efficiency [21]. Despite their marked obesity, however, LFABP^-/-^ mice were metabolically healthy, being normoglycemic, normoinsulinemic, and normolipidemic, and displayed a protection against hepatic steatosis [21,49]. Additionally, HF fed LFABP^-/-^ mice were more active and had greater exercise endurance than WT mice [21,27]. Since LFABP is highly expressed in both the liver and the intestine, we sought to determine whether its ablation in either of these two tissues, or both, underlies the MHO phenotype found in the whole body LFABP^-/-^mouse model. In this report we present data for female mice from the two conditional knockout strains relative to the LFABP^fl/fl^ controls.

Similar to the whole-body LFABP^-/-^ mice, the female LFABP^liv-/-^ and LFABP^int-/-^ mice displayed an obese phenotype when fed a HFD, with higher body weight and FM percentage when compared with LFABP^fl/fl^ counterparts. The changes observed were significantly different from similarly fed LFABP^fl/fl^, but each were of smaller magnitude than what we had observed in the whole body LFABP null mice [21]. We did not observe significant changes in energy intake, feeding efficiency, intestinal motility and fecal output, or EE. Nevertheless, the data point to a contribution from both intestinal LFABP and liver LFABP to the whole-body MHO phenotype, with smaller physiological changes in each cKO than in the whole body LFABP knockout mouse.

This dual contribution is also evidenced by the higher endurance exercise capacity observed in both cKO lines relative to control mice, suggesting that the ablation of LFABP specifically in the liver or the intestine is able to influence exercise activity and induce the same retained exercise capacity displayed by HF fed whole-body LFABP^-/-^ mice [27]. Interestingly, we recently found that whole-body LFABP^-/-^ mice have higher fecal levels of bacterial SCFA metabolites including acetate, propionate, butyrate, and other SCFAs [22]. Many studies have focused on inter-organ crosstalk between the gut and skeletal muscle via the proposed “gut-muscle axis”, highlighting the beneficial effects of SCFAs in increasing the availability of muscle glycogen and stimulating FA uptake and oxidation, resulting in more efficient energy utilization and promoting higher exercise activity [27,50–52]. It is possible that the liver- or the intestine-specific ablation of LFABP is sufficient to induce the exercise phenotype via communication between skeletal muscle, intestine, and liver, via alteration in microbiome composition and SCFA levels. An analysis of fecal microbiota in the conditional knockout mice is currently underway.

LFABP bind FAs as well as other hydrophobic ligands including monoacylglycerol-and ethanolamide-based endocannabinoids [13,19,20]. Thus, we hypothesized that ablation of LFABP specifically in the liver would disturb FA uptake and metabolism. Indeed, protection against hepatic lipid accumulation and steatosis in response to HF feeding has been shown upon whole body LFABP ablation [25,49,53]. Here we found that the liver weight to body weight ratio of LFABP^liv-/-^ mice was significantly reduced relative to the LFABP^fl/fl^ control mice, indicating a possible protection against hepatic steatosis. Moreover, while it is commonly found that obesity is associated with metabolic fatty liver disease [54–56], LFABP^liv-/-^ mice displayed no increased accumulation of neutral lipids including TG, CE, and their FA precursors, when compared with controls, despite the obese phenotype. These lower liver lipid levels were not due to alterations in VLDL secretion in the LFABP^liv-/-^ mice, nor were they due to an increase in liver tissue FA oxidation, in agreement with Erol *et al.* who also showed no reduction in hepatic FA oxidation in liver homogenates of whole body LFABP^-/-^ mice [57]. The underlying cause of reduced liver lipids in the LFABP^liv-/-^ mice is likely reduced tissue uptake; FA uptake experiments, which measured the activity of gavaged ^14^C-oleate in different tissues, demonstrated a significant reduction in FA uptake in the liver of LFABP^liv-/-^ mice relative to the floxed control mice. The reduced hepatic FA uptake observed here is in agreement with prior literature where the functions of LFABP were examined in vitro in cultured transformed cells, cultured primary hepatocytes from LFABP^-/-^ mice, and in the null mice themselves; a reduction in hepatic FA uptake was also suggested to account for the observed protection against hepatic steatosis in HF fed whole-body LFABP^-/-^ mice [49,53]. Reduced FA uptake into liver indicates the potential of increased FA availability to other tissues. A trend toward higher FA uptake was found in LFABP^liv-/-^ adipose tissue when compared to control mice, suggesting that more FA are taken up by adipose tissue for storage as TG; this may, at least in part, explain the greater FM seen in the LFABP^liv-/-^ mice. We also explored whether the observed protection against hepatic steatosis in the LFABP^liv-/-^ mice might also be influenced by compensatory responses in the intestine to the liver-specific ablation of LFABP, since LFABP is still expressed in the proximal small intestine where intestinal lipid processing primarily occurs. The intestinal mucosa of the LFABP^liv-/-^ mice showed higher TG levels than the control LFABP^fl/fl^ mice. We found previously that intestinal-LFABP plays a role in incorporation of FA into TG [21,36], thus the presence of LFABP in the intestine along with possibly increased FA availability, could underlie the increased intestinal mucosal TG levels. Additionally, it has been shown that obesity is associated with a substantial suppression in rates of intestinal TG secretion [26,58]; here the obese LFABP^liv-/-^ mice showed a decrease in the rate of TG appearance in the blood after a lipid bolus, relative to floxed controls. This reduction in chylomicron secretion rates could also contribute to the higher intestinal TG level. In this regard, it is of interest that the LFABP^int-/-^ mice had reduced net FA intestinal uptake and did not maintain control levels of TG secretion, rather they also showed a significant reduction in chylomicron secretion rates. Taken together, these finding support the role of intestinal LFABP in chylomicron formation and assembly, likely via its role in the generation of pre-chylomicron transport vesicles[59,60].

It has been hypothesized that LFABP acts to carry ligands to the lipophilic binding pockets of nuclear hormone receptors (NHRs) such as peroxisome proliferator-activated receptor alpha (PPARα), potentially allowing for LFABP to influence the regulation of lipid metabolism-related genes [61–63]. While LFABP may affect trafficking of FA to the nucleus via its interactions with PPARα, LFABP is nevertheless not required for the action of PPARα [57,64]. Here we found that liver-specific ablation of LFABP altered expression of hepatic lipid metabolic pathways, including those involved with FA transport, FA synthesis, and FA oxidation. However, as noted, these transcriptional changes did not lead to discernable effect on liver lipids levels. By binding FAs, LFABP functions to generate a FA concentration gradient across the plasma membrane, promoting FA uptake upon binding and transporting FA into various metabolic pathways [65,66]. Ablation of liver-LFABP likely leads to more unbound FA in the cytosol, hence lowering the concentration gradient to limit further FA uptake. Therefore, the reduction in the expression of some genes involved in FA uptake and synthesis could be a negative feedback mechanism to prevent accumulation of unbound FA. The imbalanced FA concentration that might be caused by decreased uptake and synthesis of FA was also restored by reducing FA degradation, as suggested by decreased expression of genes involved in FA oxidation. Overall, these changes appear to lead to balanced hepatic FA and TG concentrations that were comparable to those found in LFABP^fl/fl^ control mice. Taken together, the gene expression data support the pivotal role of LFABP in hepatic FA uptake and trafficking. Notably, the ablation of liver-LFABP did not appreciably affect intestinal gene expression. By contrast there was a substantial decrease in the expression of many hepatic genes involved in FA metabolic pathways upon ablation of intestinal-LFABP. This suggests crosstalk between the intestine and the liver, and requires further investigation.

In summary, hepatic- or intestinal-LFABP ablation in female mice appears sufficient to partially induce the MHO phenotype of the whole body LFABP^-/-^ mice. The tissue-specific ablation of LFABP in liver or intestine resulted in increased body weight and FM relative to floxed control mice, although the extent of these phenotypic changes was not as dramatic as those observed in whole body LFABP^-/-^ mice [21,23,35]. Despite their obese phenotype, both LFABP^liv-/-^ and LFABP^liv-/-^ cKO mice were protected against a HFD induced decline in endurance activity. In addition, the obese LFABP^liv-/-^ mice were protected against the development of hepatic steatosis, with the reduced neutral lipid accumulation relative to HFD-fed control mice likely due to the reduction in hepatic FA uptake, with FA redistribution to adipose tissue.

It is noteworthy that the inhibition of various members of the FABP family, either genetically or pharmacologically, is often associated with positive health outcomes. For example, FABP7 and FABP5 inhibitors have been reported to ameliorate symptoms of multiple sclerosis in a mouse model [67]; macrophage-specific ablation of FABP4 or FABP5 in mice is protective against atherosclerosis [68]; an FABP3 inhibitor was found to prevent the α-synuclein toxicity that appears to play a role in neurological disorders such as Parkinson’s disease and dementia [69]; and numerous cancers show elevated levels of FABPs, with gene suppression or ablation inhibiting tumor progression [70]. The present findings suggest that development of an FABP1/LFABP inhibitor might prove useful in preventing both obesity-associated liver steatosis and the decline in exercise performance.

## Author Contributions

Conceptualization, HRT, AIL, LQ and JS; formal analysis, HRT and AIL; investigation, HRT, AIL, AD, YZ, SZ, YHL, and JS; resources, JS; data curation, HRT, AIL, JS; writing—original draft preparation, HRT, AIL, and JS; writing—review and editing, HRT, AIL, JS; supervision, JS; project administration, JS; funding acquisition, JS. All authors have read and agreed to the published version of the manuscript.

## Institutional Review Board Statement

This study was conducted in accordance with the Declaration of Helsinki and approved by the Rutgers University Institutional Review Board.

## Data Availability Statement

The raw data supporting the conclusions of this article can be made available upon reasonable request.

## Acknowledgments

We would like to thank Drs. Ghassan Yehia and Peter Romanienko from the Rutgers Genome Editing Core Facility, whose expertise was integral for generating the LFABP-cKO mice. We also acknowledge Dr. Tracy Anthony for generously providing the Albumin-Cre mice.

