## Supplemental information for "Tissue-Specific Ablation of Liver Fatty Acid-Binding Protein Induces a Metabolically Healthy Obese Phenotype in Female Mice"

**Supplemental Material**

*Generation of LFABP Floxed Mice*

Two separate single guide RNAs (sgRNAs) were developed, with each formed to contain a targeting sequence (crRNA), and a Cas9 nuclease-recruiting sequence (tracrRNA) [71]. The crRNA is a 20-nucleotide sequence that is homologous to sequences either upstream or downstream of exons 2 and 3 respectively of the *Fabp1* gene. This sequence is adjacent to a sequence called a protospacer adjacent motif (PAM), which is required for Cas9 recognition, directing Cas9 nuclease activity specifically to the *Fabp1* alleles. The Cas9 nuclease induces double-strand breaks (DSBs) at the sites of recognition. To allow for a precise editing to occur, single stranded (ss) DNA donors were used to introduce 2 loxP sites within the host DNA via the homology directed repair mechanism. The ssDNA sequence is the same as for the WT *Fabp1* gene and contains the crRNA targeting sequence. Once the loxP site was successfully added, the crRNA sequence was separated from the PAM sequence, to prevent further DSBs from occurring again. The ssDNA donor that is added upstream to exon 2 of *Fabp1* contains also a PsiI restriction site, while the ssDNA donor that is added downstream to exon 3 contains an EcoRI restriction site (**Supplemental Figure 1**).

The ssDNA donors containing loxP, Cas9 protein, and the gRNA targeting downstream of the *Fabp1* gene were microinjected into the oocytes of C57BL6/N mouse. After fertilization with sperm from C57BL6/J mice, the resultant mice contained only downstream loxP site. Then the oocytes from the resultant mice were microinjected again, but this time the gRNA targeted the upstream site of the *Fabp1* gene. In this case, both exons 2 and 3 of the *Fabp1* gene would be flanked by loxP sites. The resultant floxed LFABP (LFABP^fl/fl^) mice on the mixed C57BL6/J and C57BL6/N background were backcrossed with WT C57BL6/J mice 4 additional times, yielding congenic LFABP^fl/fl^ mice on the C57BL6/J background.

**Supplemental Figure 1.** Sequences for the upstream and downstream ssDNA donors

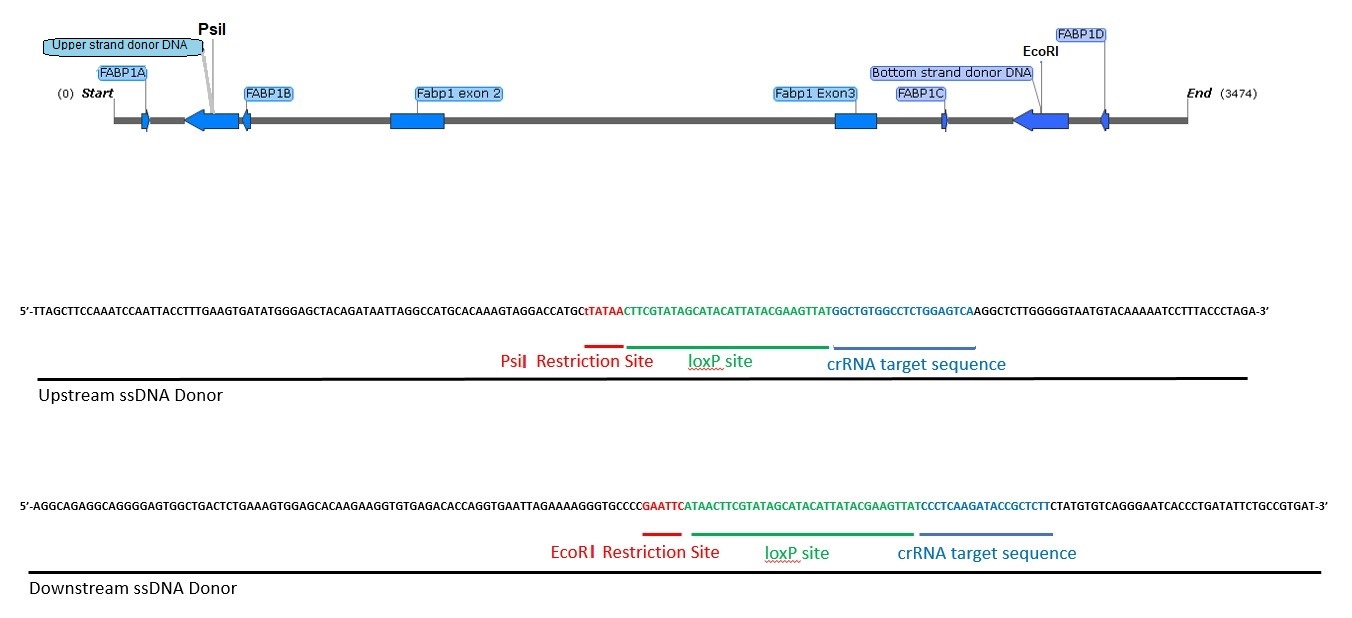

The upstream ssDNA donor contains a PsiI restriction site, loxP sequences, and distinct crRNA targeting sequences, while the downstream ssDNA donor contains an EcoRI restriction site, loxP sequences, and distinct crRNA targeting sequences. PsiI and EcoRI restriction sites are important for assessing the presence of both the upstream and downstream loxP sites.

**Supplemental Table 1.** Primers sequences for qPCR analysis of lipid metabolism

| **Genes** |  | **Sequences (5’🡪3’)** |
| --- | --- | --- |
| ***Tbp*** | Forward | 5′-AGAACAATCCAGACTAGCAGCA-3′ |
|  | Reverse | 5′-GGGAACTTCACATCACAGCTC-3′ |
| ***Fatp2*** | Forward | 5′-TCCTCCAAGATGTGCGGTACT-3′ |
|  | Reverse | 5′-TAGGTGAGCGTCTCGTCTCG-3′ |
| ***Fapt5*** | Forward | 5′-TCTATGGCCTAAAGTTCAGGCG-3′ |
|  | Reverse | 5′-CTTGCCGCTCTAAAGCATCC-3′ |
| ***Got1*** | Forward | 5′-GCGCCTCCATCAGTCTTTG-3′ |
|  | Reverse | 5′-ATTCATCTGTGCGGTACGCTC-3′ |
| ***CD36*** | Forward | 5′-ATGGGCTGTGATCGGAACTG-3′ |
|  | Reverse | 5′-GTCTTCCCAATAAGCATGTCTCC-3′ |
| ***Ldlr*** | Forward | 5′-TGACTCAGACGAACAAGGCTG-3′ |
|  | Reverse | 5′-ATCTAGGCAATCTCGGTCTCC-3′ |
| ***Lfabp*** | Forward | 5′-ATGAACTTCTCCGGCAAGTACC-3′ |
|  | Reverse | 5′-CTGACACCCCCTTGATGTCC-3′ |
| ***Scp2*** | Forward | 5′-CCTTCTGTCGCTTTGAAATCTCC-3′ |
|  | Reverse | 5′-GCTTCCTTTGCCATATCAGGAT-3′ |
| ***Acbp*** | Forward | 5′-GAATTTGACAAAGCCGCTGAG-3′ |
|  | Reverse | 5′-CCCACAGTAGCTTGTTTGAAGTG-3′ |
| ***Acc1*** | Forward | 5′-ATGGGCGGAATGGTCTCTTTC-3′ |
|  | Reverse | 5′-TGGGGACCTTGTCTTCATCAT-3′ |
| ***Fasn*** | Forward | 5′-GGAGGTGGTGATAGCCGGTAT-3′ |
|  | Reverse | 5′-TGGGTAATCCATAGAGCCCAG-3′ |
| ***Elovl6*** | Forward | 5′-GCACCCGAACTAGGTGACAC-3′ |
|  | Reverse | 5′-CCCCAGCGACCATGTCTTT-3′ |
| ***Scd1*** | Forward | 5′-TTCTTGCGATACACTCTGGTGC-3′ |
|  | Reverse | 5′-CGGGATTGAATGTTCTTGTCGT-3′ |
| ***Acsl1*** | Forward | 5′-TGCCAGAGCTGATTGACATTC-3′ |
|  | Reverse | 5′-GGCATACCAGAAGGTGGTGAG-3′ |
| ***Acsl5*** | Forward | 5′-TCCTGACGTTTGGAACGGC-3′ |
|  | Reverse | 5′-CTCCCTCAATCCCCACAGAC-3′ |
| ***Cpt1α*** | Forward | 5′-CTCCGCCTGAGCCATGAAG-3′ |
|  | Reverse | 5′-CACCAGTGATGATGCCATTCT-3′ |
| ***Cpt2*** | Forward | 5′-CAGCACAGCATCGTACCCA-3′ |
|  | Reverse | 5′-TCCCAATGCCGTTCTCAAAAT-3′ |
| ***Acadl*** | Forward | 5′-GAGAAGTGAGTAGAGAGGTCTGG-3′ |
|  | Reverse | 5′-AACTGCTGTTGAGAGCAAGTC-3′ |
| ***Hmgcs2*** | Forward | 5′-AGAGAGCGATGCAGGAAACTT-3′ |
|  | Reverse | 5′-AAGGATGCCCACATCTTTTGG-3′ |
| ***Acox1*** | Forward | 5′-TAACTTCCTCACTCGAAGCCA-3′ |
|  | Reverse | 5′-AGTTCCATGACCCATCTCTGTC-3′ |
| ***Acaa1b*** | Forward | 5′-TGCAGTCAAGCACAAGCCT-3′ |
|  | Reverse | 5′-CAGGGAGTTCAGGGTGCTAC-3′ |
| ***Pparα*** | Forward | 5′-AGAGCCCCATCTGTCCTCTC-3′ |
|  | Reverse | 5′-ACTGGTAGTCTGCAAAACCAAA-3′ |
| ***Gpat1*** | Forward | 5′-CTTGGCCGATGTAAACACACC-3′ |
|  | Reverse | 5′-CTTCCGGCTCATAAGGCTCTC-3′ |
| ***Lipc*** | Forward | 5′-ATGGGAAATCCCCTCCAAATCT-3′ |
|  | Reverse | 5′-GTGCTGAGGTCTGAGACGA-3′ |
| ***Hnf1α*** | Forward | 5′-GACCTGACCGAGTTGCCTAAT-3′ |
|  | Reverse | 5′-CCGGCTCTTTCAGAATGGGT-3′ |
| ***Hnf4α*** | Forward | 5′-CACGCGGAGGTCAAGCTAC-3′ |
|  | Reverse | 5′-CCCAGAGATGGGAGAGGTGAT-3′ |
| ***Fgf21*** | Forward | 5′-AGATCAGGGAGGATGGAACA-3′ |
|  | Reverse | 5′-TCAAAGTGAGGCGATCCATA-3′ |
| ***Gpat3*** | Forward | 5′-GGCCTTCGGATTATCCCTGG-3′ |
|  | Reverse | 5′-CTTGGGGGCTCCTTTCTGAA-3′ |
| ***Ucp2*** | Forward | 5′-TAGTGCGCACCGCAGCC-3′ |
|  | Reverse | 5′-AGCTCATCTGGCGCTGCAG-3′ |
| ***Mttp*** | Forward | 5′-AGCCAGTGGGCATAGAAAATC-3′ |
|  | Reverse | 5′-GGTCACTTTACAATCCCCAGAG-3′ |

Acaa1b: Acetyl-CoA acyl transferase 1b; Acbp: Acyl-CoA-binding protein; Acc1: Acetyl-CoA carboxylase 1; Acox1: Acyl-CoA oxidase-1; Acsl: Acyl-CoA synthetase long chain; CD36: Cluster of differentiation 36; Cpt: Carnitine palmitoyl transferase; Elovl6: Long chain fatty acyl elongase; Fasn: Fatty acid synthase; Fatp: Fatty acid transport protein; Fgf21: Fibroblast growth factor 21; Got, Glutamic oxaloacetic transaminase; Gpat, Glycerol-3-phosphate acyltransferase; Hmgcs: 3-hydroxy-3-methylglutaryl-CoA synthase; Hnf: Hepatocyte nuclear factor; Lfabp; Liver fatty acid-binding proteins; Lipc: Hepatic TG lipase; Ldlr, Low-density lipoprotein receptor; Mttp: Microsomal triglyceride transfer protein; Ppar α, Peroxisome proliferator-activated receptor alpha Scp2, Sterol carrier protein2; Tbp: TATA-binding protein; Ucp2, mitochondrial uncoupling protein2.
